# A target product profile for a rapid diagnostic test to monitor mosquito gene drive presence and frequency

**DOI:** 10.64898/2025.12.16.694691

**Authors:** Prateek Verma, Sebald Verkuijl, Calvin K. Yee, Filippo Randazzo, Damaris Matoke-Muhia, Jonathan Kayondo, Nikolai Windbichler, Michael R. Santos, Frédéric Tripet, John M. Marshall

**Affiliations:** Divisions of Biostatistics & Epidemiology, School of Public Health, University of California, Berkeley, California, United States of America; Department of Life Sciences, Faculty of Natural Sciences, Imperial College London, London, United Kingdom; Leverage Science, LLC, Berkeley, California, United States of America; Centre for Biotechnology Research and Development, Kenya Medical Research Institute, Nairobi, Kenya; Department of Entomology, Uganda Virus Research Institute, Uganda; Foundation for the National Institutes of Health, North Bethesda, Maryland, United States of America; Swiss Tropical and Public Health Institute, Kreuzstrasse 2, Allschwil, Switzerland; University of Basel, Petersplatz 1, Basel, Switzerland

**Keywords:** *Anopheles* mosquitoes, gene drive, malaria control, mosquito monitoring, pooled samples, rapid diagnostic test, target product profile

## Abstract

Malaria remains a major global health challenge, with over 263 million cases and nearly 600,000 deaths reported in 2023, the majority in sub-Saharan Africa. While conventional interventions such as insecticide-treated nets, indoor residual spraying and antimalarial drugs have reduced transmission, progress has stalled due to the limitations of these interventions and the emergence of resistance. Gene drive-modified mosquitoes represent a promising, potentially transformative vector control strategy, capable of spreading malaria-refractory traits or suppressing mosquito populations. Successful field deployment will depend upon monitoring systems to track the presence and frequency of gene drive constructs as they spread and persist. Current molecular surveillance techniques, though effective, are resource-intensive and reliant on laboratory infrastructure and technical competencies. Here, we make the case for a near-universal and low-cost rapid diagnostic test (RDT) designed to detect gene drive mosquitoes in the field, to complement existing surveillance infrastructure. Two use cases are outlined: i) to detect the presence of the drive construct in a new population, and ii) to provide an estimate of drive frequency prior to more accurate laboratory-based measurements. We provide a target product profile for the RDT outlining minimally essential and ideal characteristics, including test procedures, sensitivity, specificity, usability by a range of stakeholders in field settings, and compatibility with pooled testing of mosquito samples. An RDT for gene drive construct detection would support community access and participation in monitoring, enhance regulatory oversight, and promote transparency in field trials, thereby facilitating responsible deployment of gene drive-based malaria interventions.

## 1. Background

Malaria has caused the deaths of tens of millions of people over the past 50 years, including an estimated 263 million cases and 597 thousand deaths worldwide in 2023 [1]. Exacerbated by poverty and limited economic development, over 95% of recent cases and deaths have occurred in sub-Saharan Africa. Substantial reductions in malaria transmission have been achieved since 2000 with the widespread adoption of interventions including long-lasting insecticide-treated nets (LLINs), indoor residual spraying with insecticides (IRS), and artemisinin combination therapy drugs (ACTs) [2]; however, progress has stalled over the last decade due to limitations of these interventions, emergence of resistance, and financial constraints [1]. Novel malaria control tools are urgently needed, and gene drive-modified mosquitoes are one of the most promising families of new control tools currently in the development pipeline. Gene drive is a phenomenon whereby “a particular heritable element biases inheritance in its favor, resulting in it being found in most progeny, and over successive generations becoming prevalent in the whole population” [3]. Gene drive developers are currently focusing on “low-threshold” approaches for targeting the primary malaria vectors in Africa. These systems require the release of only small numbers of modified mosquitoes in order for the construct to spread through a population, thus having a high impact on malaria transmission at relatively low production and delivery costs. Antimalarial effects can be achieved through suppressing mosquito population size [4], or modifying mosquitoes to be less capable parasite vectors [5], or a combination of the two [6].

Efficient population suppression has been demonstrated in small and large cage experiments by targeting female reproduction in *Anopheles gambiae* [4,7,8]. Additionally, antimalarial population modification strategies have been demonstrated with modifications such as single-chain variable-fragment monoclonal antibodies targeting ookinetes and sporozoites in *Anopheles stephensi, An. gambiae* and *Anopheles coluzzii* [5,9,10], and midgut-expressed antimicrobial peptides affecting the development of malaria sporozoites in *An. gambiae* [11]. Mathematical modeling suggests that both suppression and modification strategies could greatly reduce malaria transmission in the field [5,12,13]; however, the success of these strategies largely depends upon parameters that can only be reliably measured in the field [14]. Field trials are therefore essential prior to widespread use, and will require acceptance from many stakeholders, including affected communities, local authorities, National Malaria Control Programs (NMCPs)/National Malaria Elimination Programs (NMEPs), governments and regulatory agencies.

Monitoring of the first gene drive field trials will be particularly important, given the novelty of the intervention, the degree of interest surrounding it, and the potential for field trials to proceed in an adaptive manner, based on data collected as the intervention proceeds [15]. Interest in the intervention provides a strong motivation to include community members in monitoring activities, ideally facilitated by established entities such as schools and NMCPs. A robust monitoring protocol will also be needed to adapt trial design in response to data collected during the study. This is particularly important given uncertainties in intervention parameters that will only be resolved during or following a release. For instance, mathematical models to support trial design will initially be based on laboratory data pertaining to the gene drive construct, and ecological data pertaining to the target mosquito species; but the fitness of introgressed gene drive mosquitoes can only be reliably measured in the field, and is expected to be highly influential on the optimal release scheme [16]. When designing a monitoring program, careful consideration should be given to data requirements to support trial objectives, as well as to optimal sampling schemes, mosquito capture techniques, and molecular assays to support these [17]. The nature of the gene drive construct is also relevant, as a suppression gene drive may only have a transient presence at a location before locally removing the target species (and itself), if successful. Following a field trial, intermediate to long-term monitoring may be required by regulators, including at areas beyond the release site.

In the event of approval for widespread deployment as a novel malaria control intervention, monitoring activities can be calibrated to more closely match the needs of malaria control programs in terms of scalable logistics and costs. There is a need for novel technologies and approaches to address the complex monitoring needs of gene drive field trials and post-release monitoring. Here, we argue that the process of gaining public and regulatory acceptance for gene drive field testing would greatly benefit from the development of a simple, low-cost, rapid diagnostic test (RDT), usable by specialists and non-specialists, to distinguish gene drive from non-gene drive mosquitoes in any setting. Such a tool would support regulators and biosafety agencies, and help build trust by empowering local communities to participate in monitoring efforts. The RDTs could also be shared with NMCPs, vector-borne disease control units, and other stakeholders and release programs, thereby enhancing transparency and fostering collaboration. RDTs could also be shared with researchers conducting field trials of other malaria control tools in nearby areas. Accuracy of information collected from these tests will be of key importance, and hence ideal partners in RDT use will have an established presence in the surveillance area, and existing mechanisms for entomological collections, quality control and accountability. This will require an operational plan for implementation, including workflows, training, data management, and adequate funding to support these activities.

### Box 1.

**Current mosquito sampling and genotyping strategies**

Depending on location, a monitoring program that samples mosquitoes monthly can collect anywhere from a few mosquitoes per village per week in the dry season to several hundred or thousands per village per week at the peak of the rainy season(s) [18,19]. Common adult sampling techniques include pyrethroid spray catches (PSCs), indoor aspirations, and/or the use of traps [20]. Common species are usually identified taxonomically, and target species are stored either in ethanol 75-80% or dried with silica gel ahead of molecular analyses used to characterize morphologically identical members of the *An. gambiae* complex or *An. funestus* group. Once in the laboratory, DNA is extracted from samples using homogenization followed by either cheap extractions based on either buffer digestion and ethanol precipitation with differential centrifugation, or a more expensive affinity column-based extraction. The former approach is used for processing large volumes at lower cost using simple PCR with fragment visualization on agarose gels [21,22], while the latter approach is used when downstream analyses require purer DNA, such as multiplexed or nested PCR, qPCR, and sequencing-based analyses, or when the target DNA is rare compared to mosquito genomic DNA (e.g., mosquito pathogen detection) [23,24].

Most African institutions leading and contributing to mosquito surveillance programs, and/or participating in the baseline studies informing gene drive mosquito release programs, have existing workflows for individual mosquito sample processing based around simple PCR-based approaches for species diagnostics, and sometimes bloodmeal analyses, whilst *Plasmodium* detection can be performed either by nested PCR, Taqman qPCR, or ELISA. Hence, in the context of gene drive field trial monitoring, assays for the genetic construct should fit into existing workstreams based on PCR and make use of existing equipment that does not require novel or complex supply chains.

For simple genetic construct detection assays, PCR can provide a presence/absence signal if primers are designed to bind to sequences only found on the transgene. Through adding a primer specific to the wild-type sequence, the resulting three-primer multiplexed PCR assays can be used to discriminate between heterozygotes and homozygotes [25]. Care must be taken to design primers that consider local sequence polymorphisms, as mutations in priming regions are the most common issue affecting the specificity of PCR-based approaches used on field-collected samples [22,26]. When mosquito samples are abundant, or for large-scale monitoring, genetic construct detection assays that interrogate pools of mosquitoes may become cost-effective. However, detecting rare genetically modified mosquitoes in large pools may exceed the sensitivity of PCR, which cannot reliably detect targets below 1:100. For this, other molecular assays such as loop-mediated isothermal amplification (LAMP) may be better suited. Like PCR, LAMP amplifies specific DNA regions, but operates at a constant 60-65°C temperature, eliminating the need for a thermocycler and enabling faster turnaround, as LAMP products can be detected using fluorescence with intercalating dyes, or colorimetric methods [27,28]. LAMP requires strict standard operating procedures (SOPs), dedicated instruments, and workspace specifications, as its high sensitivity makes it prone to contamination. While LAMP assays can be multiplexed, it is more challenging than with PCR since each target requires at least four primers [29,30]. As molecular technologies advance and institutions in malaria-endemic countries gain better access to funding and equipment, new cost-effective genotyping options for monitoring genetic constructs may become available; however, high-tech solutions are not always the most appropriate, as they require dedicated facilities and trained personnel.

## 2. Use cases for a gene drive rapid diagnostic test

During field releases, gene drive mosquitoes are expected to, over successive generations, disperse outwards from points of release, a process that may be influenced by environmental factors such as wind and barriers to movement. Ideally, local mosquito dispersal patterns will be empirically quantified through mark-release-recapture (MRR) studies prior to gene drive release. These pre-release studies can inform the spatial delineation of post-release surveillance zones for monitoring gene drive mosquito presence and frequency [31]. However, mosquito dispersal is frequently characterized as long-tailed, with occasional long-distance movements occurring beyond the predicted range [32], possibly facilitated by anthropogenic factors such as transportation or naturally by wind [33]. In this context, there is a need for a gene drive rapid diagnostic test (RDT) - a tool that can function independently of the powerful, but likely inflexible, core field trial monitoring systems (**Box 1**). We propose two specific RDT use cases illustrated schematically in **Fig. 1**: i) detection of the gene drive construct from pools of mosquitoes caught at release and post-release surveillance zones as well as distant areas where the construct is not expected to have reached, and ii) approximate quantification of a gene drive “carrier frequency” (the proportion of a population either heterozygous or homozygous for the drive) through the use of multiple tests of pooled samples in areas where the gene drive has been confirmed present.

**Fig. 1.**
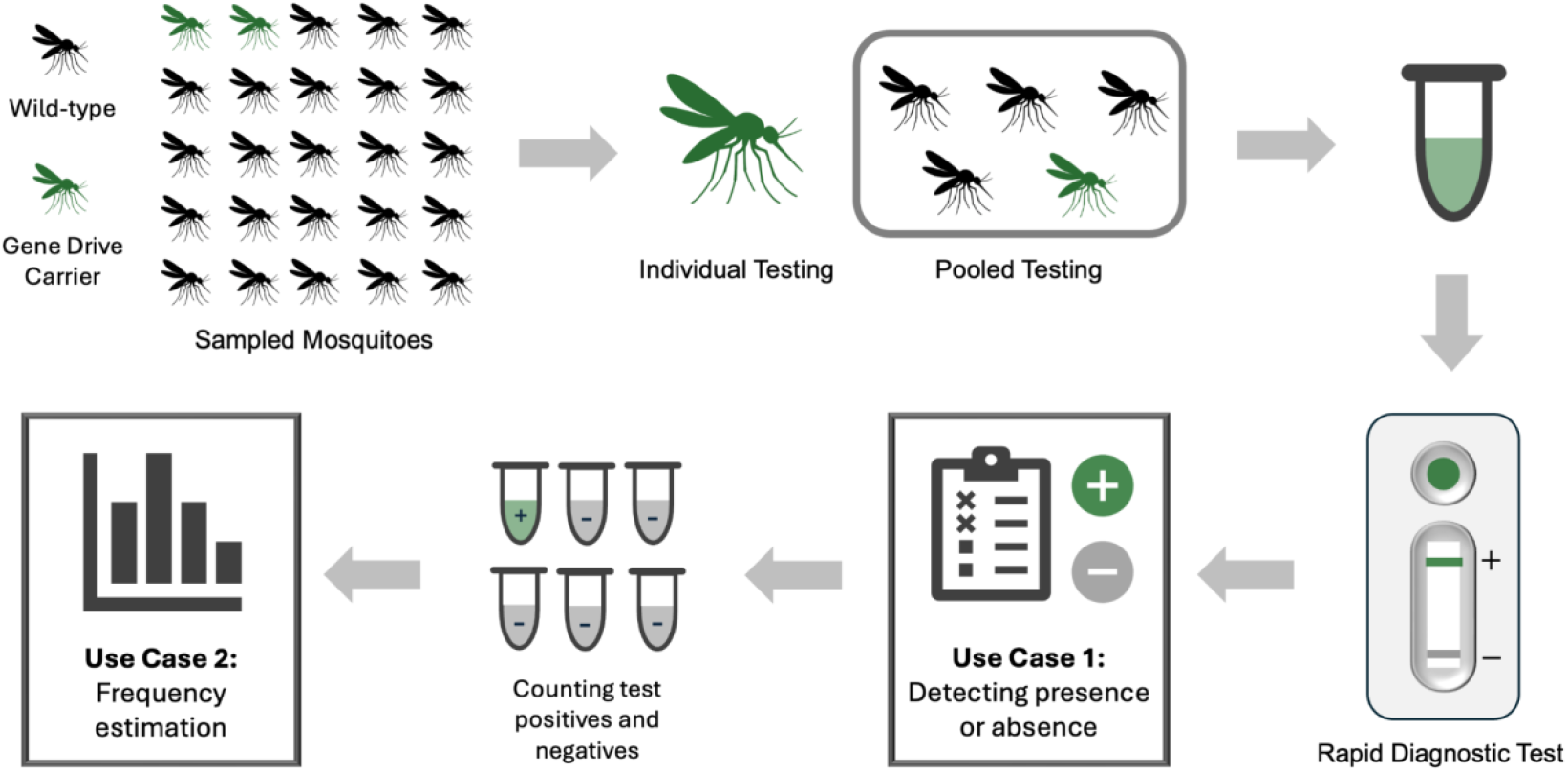
Schematic workflow for performing RDTs on sampled mosquitoes for gene drive monitoring. Mosquitoes can be tested individually or in pools for the two use cases. Use case 1 involves detecting the presence or absence of gene drives, while use case 2 estimates the frequency of gene drive carriers through counts of positive and negative tests.

### 2.1. Use case 1: Detecting the presence of gene drive mosquitoes

This use case focuses on the timely detection of the gene drive construct in a mosquito population at a time when its presence has not yet been confirmed. The RDT would serve as a complement to core monitoring activities of field trials, and as a low-cost tool to enable easy participation of communities in trial and monitoring areas, as well as engagement of specific stakeholder groups, in the context of structured and active engagement activities. Locations where the gene drive has not yet been detected could include designated release and post-release monitoring zones, as well as locations beyond the mosquito’s predicted dispersal range. We anticipate the RDT will often be used in locations where gene drive mosquitoes are expected to be absent, and are not part of the core monitoring pipeline or its buffer zones. In this context, RDTs offer distinct advantages, foremost among them the capacity to enable active participation of local communities and stakeholders in surveillance activities within defined monitoring areas. Such capacity is essential for maintaining designated control areas as true negative controls, ensuring their validity for comparative analyses and for detecting faster-than-expected spread of the gene drive. RDTs provide clear utility in regions lacking access to centralized laboratory facilities or staff trained in molecular assays. Collections would be made through breeding sites, traps placed in houses overnight, or indoor aspiration performed in a house or animal shed. If samples are found to be positive for the gene drive, these should be confirmed through additional RDT tests and/or laboratory-based assays, and these locations can then be folded into an existing surveillance pipeline. In summary, for use case 1, the RDT would: i) facilitate detection in low-resource and isolated settings, ii) enable participation of communities in trial and monitoring areas in the context of structured engagement activities, iii) provide an initial test for inclusion of a zone as part of the core monitoring system, and iv) provide an approach for detecting the gene drive after the initial trial phase.

### 2.2. Use case 2: Estimating gene drive carrier frequency

This is an extension of use case 1, and takes advantage of the simplicity and flexibility of the RDT to estimate gene drive carrier frequency by performing a limited number of RDTs on mosquito pools of various sizes. This series of RDTs can provide an estimate of gene drive carrier frequency, especially when sampled mosquitoes are representative of the population, and distributed optimally among pools. It is assumed that, when more accurate gene drive carrier or allele frequency estimates are needed, these will involve more extensive and representative sampling, and processing via molecular assays in the laboratory. The RDT-inferred carrier frequency estimates will therefore be provisional, and only seek to provide a rough estimate of gene drive carrier frequency in the population. In this context, for use case 2, the RDT would: i) facilitate estimation of gene drive carrier frequency in newly-identified positive sites, and ii) facilitate estimation of gene drive carrier frequency in known positive sites that may not be actively monitored, or would like to be audited by a third party. Collectively, these use cases place certain requirements and restrictions on the design and performance of the test.

## 3. Target product profile for a gene drive rapid diagnostic test

Here, we outline a target product profile (TPP) to guide the design of RDTs for monitoring gene drive mosquitoes, specifying the minimally essential and ideal criteria. We envision use case 1 to be the most common application of the RDT, and hence prioritize its requirements. As we discuss later, we find that the criteria for use case 1 also generally satisfy the criteria for use case 2, and hence, the minimally essential and ideal criteria for both use cases are presented jointly in **Tables 1 and 2**.

**Table 1.** Target product profile for key features of a gene drive rapid diagnostic test.

| Feature | Minimally essential | Ideal | Considerations |
| --- | --- | --- | --- |
| Target user | Specialist user with no formal training beyond a manual/leaflet | Non-specialist user with no formal training beyond a manual/leaflet | Ideally, the test would be easily adopted by parties unaffiliated with the release and monitoring efforts |
| Target use setting | Dedicated hub near the capture location | Fully on-site, at trap or larval habitat; usable handheld or on knee/ground | The test is intended for use in isolated and impromptu settings |
| Target species/scope | Validated on primary surveillance species in the target program, and no false-positives from misapplication | Valid across common vector genera (e.g., *Anopheles/Aedes*) or clearly labelled if species-restricted | Preselection of mosquito samples by species is a specialized skill |
| Analyte / target sequence(s) | Any component or product of the gene drive cassette (DNA, RNA, or protein) | Common parts across projects (e.g., Cas9 DNA/mRNA/protein; gRNA scaffold DNA/RNA) with option for construct-specific confirmatory targets | Cas9 and gRNA provide common detection targets for current and likely future implementations. If the analyte is not DNA, higher detection sensitivity will be required, as RNA and proteins will be limited to specific tissues and vary by life stage |
| Analytical sensitivity | Detect one drive-positive mosquito tested singly; detect one positive in pools of $\geq 10$ | Same as minimally essential, but with wider pool sizes (e.g., $\geq 25$) | Pool size limits would have to be balanced with reduced test cost and increased throughput |
| Sensitivity | $\geq 60\%$ (for mosquitoes tested individually), $\geq 65\%$ (for mosquitoes tested in pools of 10) | $\geq 90\%$ (for mosquitoes tested individually), $\geq 92\%$ (for mosquitoes tested in pools of 10) | Minimally essential calculations are based on $\geq 95\%$ probability of detection given a gene drive carrier frequency $\leq 5\%$ and a sample size of 100 |
| Specificity | $\geq 99.95\%$ (for mosquitoes tested individually), $\geq 99.5\%$ (for mosquitoes tested in pools of 10) | $\geq 99.98\%$ (for mosquitoes tested individually), $\geq 99.75\%$ (for mosquitoes tested in pools of 10) | Minimally essential calculations are based on a sample-wide false positive rate of $\leq 5\%$ and a sample size of 100 |
| Result type & readout | Qualitative (positive/negative), visually readable without instruments. | Quantitative/semiquantitative readout; accessible to color-vision-deficient users | Qualitative readout from pooled samples may provide an indication of allele frequency |
| Sample type | Fresh/damaged/dead adult mosquitoes; larval/pupal acceptable | Desiccated or frozen specimens | Samples may come from various sources, including traps in which mosquitoes may be dead/damaged/desiccated, or mosquitoes may be stored for some time before analyses |
| Sample collection device | Portable | Handheld | Test can be distributed and shared across sampling areas |

**Table 2.** Target product profile for test procedure and additional characteristics of a gene drive rapid diagnostic test.

| Feature | Minimally essential | Ideal | Considerations |
| --- | --- | --- | --- |
| Reagent reconstitution | All reagents supplied, with minimal reconstitution | All reagents are immediately ready to use | Access to pure water or other reagents cannot be relied upon |
| Time to result from test initiation | <120 mins | <30 mins | Tests are intended to be used on brief visits to field sites |
| Result validity stability | Up to 30 minutes | Up to 24 hours | Flexibility in readout times will enable parallel workstreams, such as setting up multiple tests simultaneously. |
| Invalid rate | ≤5% technical failure rate | ≤1% technical failure rate | Low invalid rate is required due to the relatively high cost of sample collection and brief site visits |
| **Additional characteristics** |  |  |  |
| Operating conditions | 15 to 30°C, 80% relative humidity | 15 to 30°C, 80% relative humidity | The test must work in tropical conditions |
| Test kit stability and storage conditions | No cold chain (15 to 30°C), test kit stable for at least 6 months | Same as minimally essential, but stable for at least 24 months | The test components should remain stable despite unfavorable transport and storage conditions |
| Quality control | Internal controls to confirm the validity of the test and processing | Same as minimally essential, but with each test having a unique identifier | The competence of the operator will vary, and incorrect use should be indicated; ID will ensure logged results are unique and correctly attributed |
| Remote connectivity capacity | No additional features | App-based communication to a central database with metadata such as GPS location | Data stored locally and transmitted upon obtaining cell signal. Collated test results available to relevant stakeholders |

Field compatibility and ease of use are of primary importance for the RDT, as target users ideally include non-specialists such as community members, regulators and other stakeholders, who would require no training beyond reading a manual/leaflet (**Table 1**). It is envisioned that the test would be performed nearby where mosquito collections take place - i.e., in isolated field locations far from laboratories where more in-depth analyses could take place. In the minimally essential case, the test would detect any component of the gene drive cassette, whether it be DNA, RNA or an expressed protein; but ideally, it would detect sequences that will likely be conserved between different gene drive implementations (e.g., DNA/mRNA sequences or expressed proteins related to the Cas9 component, DNA/RNA sequences related to the gRNA scaffold, or promoter DNA sequences).

For cost-effectiveness and sample processing speed, it is highly desirable that the RDT is able to detect a single positive mosquito in a pool of 10 mosquitoes (or equally, presence of the gene drive at higher positivity rates); but at a minimum, the RDT should be able to detect the gene drive construct when presented with a single gene drive-modified mosquito. Pooling mosquito samples will likely be necessary to ensure cost and throughput benchmarks can be met. There should be minimal false positives, as these would both: i) trigger costly follow-up tests, defeating the purpose of the RDT, and ii) risk generating misinformation in community engagement programs. Precise requirements for test sensitivity and specificity in the case of both individual tests and tests of pools of ten mosquitoes at a time are presented in the following section. The RDT should function on a range of mosquito sample types - at a minimum, fresh and live adults, larvae or pupae, and ideally, damaged, dead, desiccated and frozen mosquito samples as well. The RDT should be a portable, ideally handheld, device that users can implement with less than half a day of training, and ideally, without training by simply reading a manual/leaflet. Custodianship of devices and considerations regarding interpreting and sharing results should be conveyed to participants prior to RDT use, given the importance of the resulting information to community engagement programs. Interpretation of results should be visual, so that users do not require any additional instrumentation.

Minimally essential and ideal features of the test procedure are presented in **Table 2**. All reagents for the test should be supplied in a stabilized, ready-to-use format, obviating the need for specialized laboratory infrastructure. The test should ideally yield results within 30 minutes (at most two hours) as the test is intended to be used on brief visits to field sites, and the validity of the result should remain stable for at least 30 minutes, but ideally for up to 24 hours to allow time for the collation of multiple tests. The technical failure (non-result) rate for the test should be at most ≤5%, and ideally would be ≤1%. Non-results increase the required number of tests by a similar amount to the technical failure rate, which increases costs of both the test and sample collection.

Several additional test characteristics are also presented in **Table 2**. Clearly, the test must be operable in tropical environments where malaria is endemic - i.e., ~15-30°C and ~80% relative humidity. There should be no cold chain, and the test kit should be stable for at least six months. Internal controls should confirm that the test is working optimally, and invalid results should clearly be identified. It would be useful, but not essential, for the test to have the capability to connect to a phone with an application capable of sending data to a central database.

## 4. Sensitivity and specificity calculations

Here, we consider the minimally essential and ideal test sensitivity and specificity for the two use cases in more detail.

### 4.1. Test sensitivity for use case 1

In this use case, the RDT would be used to detect the presence of the gene drive cassette in a new population. The false positive rate should be low, as detection of the gene drive will likely trigger costly follow-up collections and analyses. In the minimally essential case, we would like to detect the presence of the gene drive in a population with 95% confidence when the gene drive carrier frequency reaches 5%. For a gene drive carrier frequency of *x* and a test sensitivity per mosquito of *s*, the probability of detecting the gene drive in a sample of *n* mosquitoes, *p*^*GD*^, is given by:

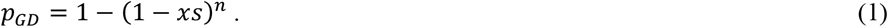

For a gene drive carrier frequency of 5% and a minimally essential probability of detection of 95%, **Fig. 2A** depicts the required sample size as a function of test sensitivity per individual mosquito. Required sample sizes are all in the vicinity of 100. Taking this as a default sample size that is achievable in the field in the rainy season, the minimally essential test sensitivity per individual mosquito is ≥60%. We then consider an ideal test sensitivity per individual mosquito of ≥90%, which leads to a probability of detection of 99% for a sample of 100 mosquitoes.

**Fig. 2.**
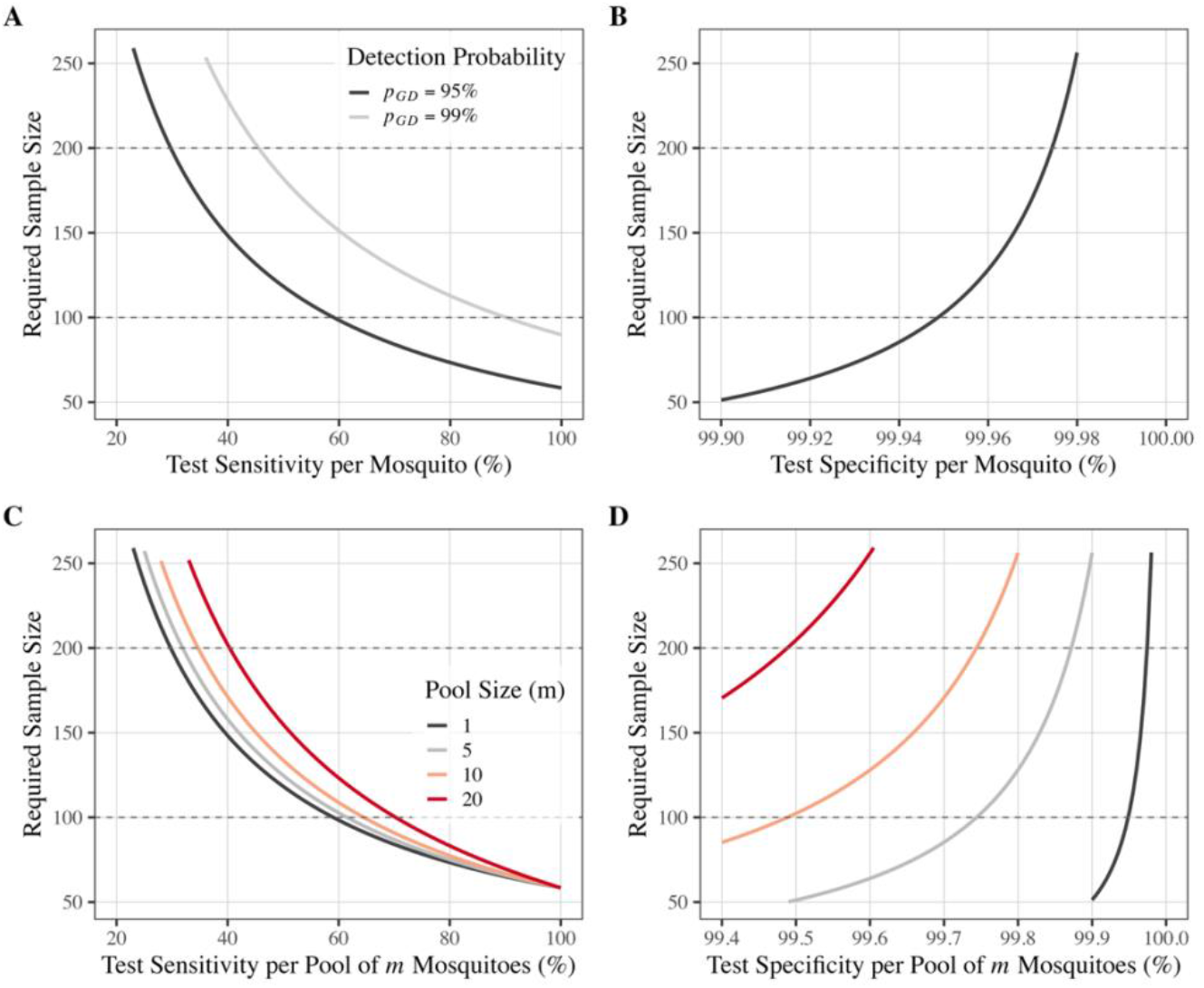
Target sensitivity and specificity parameter values for use case 1. **(A)** Required sample size as a function of test sensitivity per individual mosquito, assuming we would like to detect the presence of the gene drive in the sample with 95% (minimally essential) or 99% (ideal) confidence when the gene drive carrier frequency reaches 5%. For a sample size of 100, the minimally essential test sensitivity is ≥60% and the ideal test sensitivity is ≥90%. **(B)** Required sample size as a function of test specificity per individual mosquito, assuming a sample-wide false positive rate of 5% for a sample size of 100 (minimally essential) or 200 (ideal). **(C)** Mosquitoes are analyzed in pools of size *m*, and required sample size is depicted as a function of test sensitivity per pool (i.e., the probability of detecting at least one gene drive mosquito in a pool of this size), assuming we would like to detect the presence of the gene drive in the population with 95% confidence (minimally essential) when the gene drive carrier frequency reaches 5%. **(D)** Required sample size as a function of test specificity per pool of *m* mosquitoes (i.e., one minus the probability of detecting a false positive in a pool of this size), assuming a sample-wide false positive rate of 5% for a sample size of 100 (minimally essential) or 200 (ideal).

### 4.2. Test specificity for use case 1

Now let us consider a fully wild-type mosquito population. For a false positive rate per mosquito of *f* (corresponding to a test specificity of one minus *f*), the probability of detecting a false positive in a sample of *n* mosquitoes tested one-by-one, *p*^*FP*^, is given by:

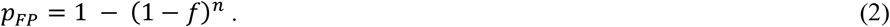

Let us continue with a total sample size of 100, per the calculations for test sensitivity. Considering a minimally essential sample-wide false positive rate of 5%, the minimally essential test specificity per mosquito is ≥99.95% (**Fig. 2B**). Increasing the sample size to 200 mosquitoes, a test specificity of ≥99.98% produces a sample-wide false positive rate of 5%. We consider this to be an ideal test specificity per mosquito. Note that, although the minimally essential test specificity of ≥99.95% for use case 1 is quite demanding, for a multiplexed diagnostic test, the composite false positive rate, *f*^*MP*^, is equal to the multiplication of false positive rates for its *M* constituent tests, i.e.:

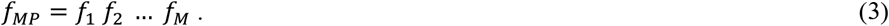

Here, *f*_1_ through *f*_*M*_ refer to the false positive rates of constituent tests 1 through *M*. I.e., for two multiplexed tests of equal specificity, the minimally essential specificity for each test is ≥97.8%, and for three multiplexed tests, the minimally essential specificity for each test is ≥92.1%.

### 4.3. Pooled analysis of samples for use case 1

Testing for the presence of a gene drive cassette in a new population will ideally be conducted with pooled samples for speed and cost-efficiency. In this case, the TPP parameters will vary depending upon the number of mosquitoes included in each pool. For a pool size of *m*, the minimally essential and ideal test sensitivity, *s*_*m*_, and test specificity, (1 - *f*_*m*_), follow a similar derivation. We first consider the test specificity for a pool of *m* mosquitoes. Given a false positive rate per pool of *m* mosquitoes, *f*_*m*_, the probability of detecting a false positive in a sample of *n* mosquitoes (where *n* is a multiple of *m*) is given by:

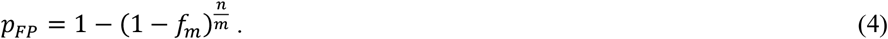

Let us consider a total sample size of 100 and a pool size of 10, as an example. Considering a minimally essential sample-wide false positive rate of <5%, the minimally essential test specificity per pool of 10 mosquitoes is ≥99.5%. If we increase the total sample size to 200 mosquitoes, a test specificity per pool of 10 mosquitoes of ≥99.75% maintains a sample-wide false positive rate of <5%. We consider this to be an ideal test specificity for a pool size of 10. Minimally essential and ideal test specificities for a range of pool sizes are depicted in **Fig. 2D**.

Next, we consider the test sensitivity for a pool of *m* mosquitoes. Again, we would like to detect the presence of the gene drive in the population with 95% confidence when the gene drive carrier frequency, *x*, reaches 5%. The chance that a gene drive carrier is present in a pool of *m* mosquitoes, *x*_*m*_, is given by:

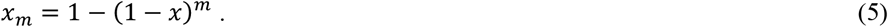

We define the test sensitivity for a pool of *m* mosquitoes, *s*_*m*_, as the probability of detecting at least one gene drive mosquito in a pool of this size. The probability of detecting a gene drive mosquito in a sample of *n* mosquitoes (where *n* is a multiple of *m*) is then given by:

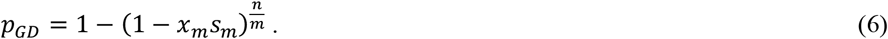

Let us consider the example of a total sample size of 100 and a pool size of 10. For a gene drive carrier frequency of 5%, the probability that a gene drive mosquito is present in a pool of 10 mosquitoes is 40.1%. Considering a minimally essential probability of detection of 95%, the minimally essential test sensitivity per pool of 10 mosquitoes is then ≥65%. We consider an ideal test sensitivity per pool of 10 mosquitoes to be ≥92%, which would lead to a probability of detection of 99% for a sample of 100 mosquitoes. Minimally essential and ideal test sensitivities for a range of pool sizes are depicted in **Fig. 2C**. Minimally essential and ideal parameter values for all scenarios explored for use case 1 are summarized in **Table 1**.

### 4.4. Test sensitivity and specificity for use case 2

In this use case, the RDT would be used to provide a ballpark estimate of gene drive carrier frequency in a population, which could be followed up with more extensive sampling and laboratory processing to provide a more accurate frequency estimate. False positives and false negatives will increase the variance in estimates of gene drive carrier frequency. They will also introduce bias into these estimates; but this can be corrected if the false positive and false negative rates are known in a field setting prior to use. For this use case, we consider two scenarios: i) measuring gene drive carrier frequency as the drive spreads through a population (e.g., for a carrier frequency of ~50%), and ii) characterizing persistence of the drive at high frequency in a population (e.g., for a carrier frequency of ~90%). Here, we consider individual tests of mosquitoes; but the case of pooled samples is described in **Additional file 1**. We are primarily interested in the standard deviation (SD) of our estimate of gene drive carrier frequency, and use a binomial distribution with a known true positive rate, *s*, and a known false positive rate, *f*, to calculate this. For a measured proportion of samples positive for the gene drive, *y*, the estimator for gene drive carrier frequency, 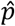, is given by:

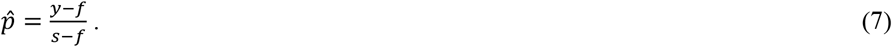

Considering a sample size, *n*, the SD of the estimator, 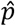, is given by:

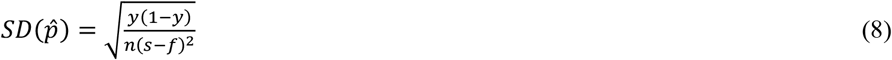

Per the previous use case, imagine a sample of 100 mosquitoes, and consider a minimally essential SD of the estimator, 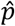, of 10%, and an ideal SD of 7% (of note, the lowest possible SD for this case is 5%, which occurs when the test sensitivity and specificity are both 100%). As the gene drive spreads through the population (i.e., for a carrier frequency of ~50%), substituting the minimally essential test sensitivity (60%) and specificity (99.95%) for use case 1 into Equation 8 yields an SD of 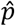 of 7.6%, and substituting the ideal test sensitivity (90%) and specificity (99.98%) for use case 1 into this equation yields an SD of 5.5%. Since these values are less than 10% and 7%, respectively, this implies that the minimally essential and ideal conditions for use case 1 automatically satisfy the conditions for use case 2 as the gene drive spreads through the population. SDs of the estimator, 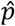, for a range of test sensitivities and specificities for this scenario are depicted in **Fig. 3A**. E.g., in the case where sensitivity is equal to specificity, the minimally essential test sensitivity/specificity is ≥75% and the ideal test sensitivity/specificity is ≥86%, and in the case where test specificity is set to 99.95%, the ideal test specificity is ≥68%.

**Fig. 3.**
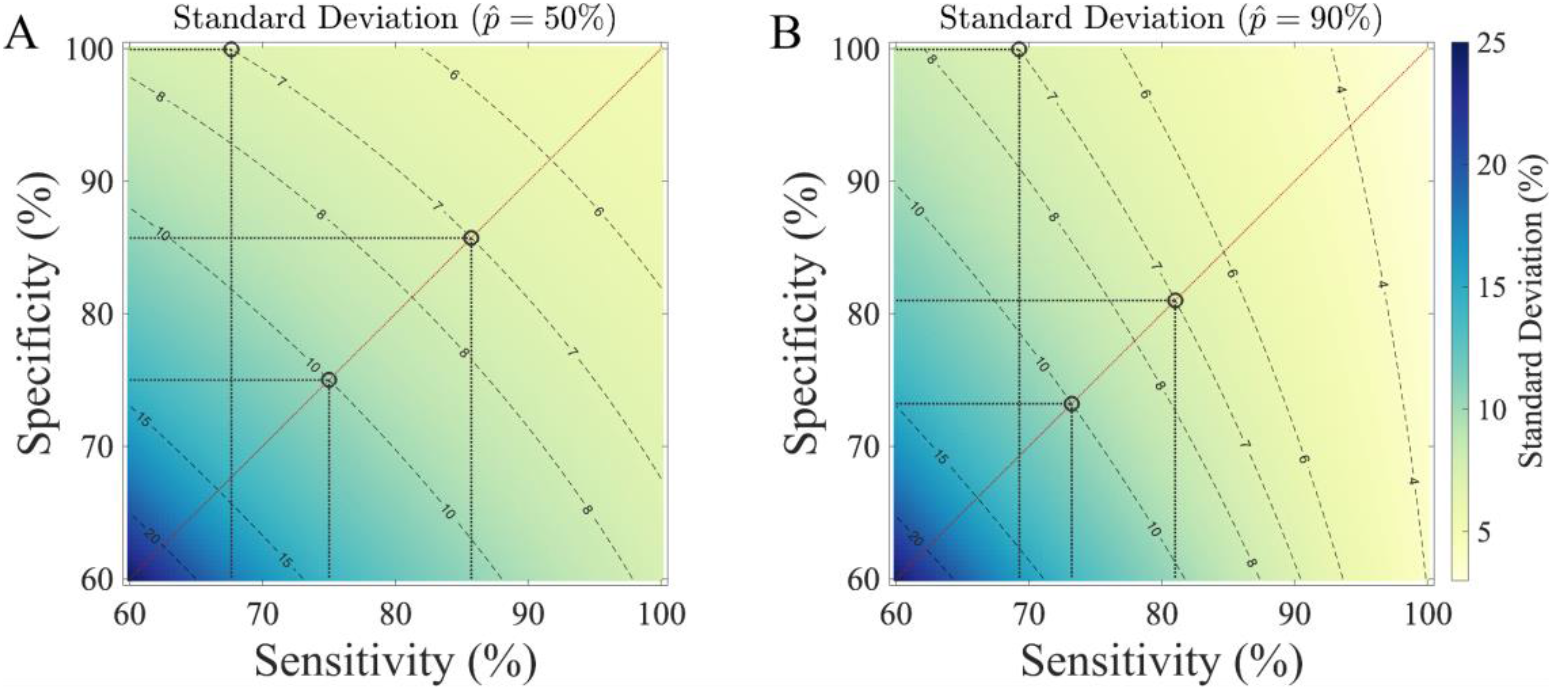
Target sensitivity and specificity parameter values for use case 2. **(A)** Standard deviation (SD) in estimated gene drive carrier frequency, 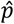, as a function of test sensitivity and specificity per mosquito as the gene drive spreads through a population (i.e., for a carrier frequency of ~50%), assuming a sample size of 100. Considering test specificity equal to sensitivity, the minimally essential test sensitivity/specificity is ≥75% (corresponding to an SD of 10%), and the ideal test sensitivity/specificity is ≥86% (corresponding to an SD of 7%). Considering a test specificity of 99.95% (for consistency with use case 1), the ideal test sensitivity is ≥68%. **(B)** SD in estimated gene drive carrier frequency as the gene drive persists in a population (i.e., for a carrier frequency of ~90%). Considering test specificity equal to sensitivity, the minimally essential test sensitivity/specificity is ≥73% (corresponding to an SD of 10%), and the ideal test sensitivity/specificity is ≥81% (corresponding to an SD of 7%). Considering a test specificity of 99.95% (for consistency with use case 1), the ideal test sensitivity is ≥69%.

Secondly, we are interested in characterizing gene drive carrier frequency as the drive persists at high frequency in a population. In this case, we consider a carrier frequency of ~90% and reapply Equation 8 for the SD of the estimator, 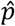. Substituting the minimally essential test sensitivity and specificity for use case 1 into this equation yields an SD of 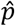 of 8.3%, and substituting the ideal test sensitivity and specificity for use case 1 yields an SD of 4.4%. Since these values are again less than 10% and 7%, respectively, this implies that the minimally essential and ideal conditions for use case 1 automatically satisfy the conditions for both scenarios for use case 2. SDs of the estimator, 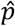, for a range of test sensitivities and specificities for the scenario where the drive persists at high frequency are depicted in **Fig. 3B**. E.g., in the case where sensitivity is equal to specificity, the minimally essential test sensitivity/specificity is ≥73% and the ideal test sensitivity/specificity is ≥81%, and in the case where test specificity is set to 99.95%, the ideal test specificity is ≥69%.

### 4.5. Pooled analysis of samples for use case 2

Given the intended use of the RDT to obtain an approximate estimate of gene drive carrier frequency in a field setting where resources are limited, it is likely that pooled rather than individual mosquito tests will commonly be used. Pooling of mosquitoes will increase the SD of the estimator,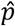; but this may be acceptable when the goal is to obtain a ballpark carrier frequency estimate that can be followed up with more extensive and rigorous molecular assays in the laboratory, when needed. For a given sample size and available number of test kits, the optimal distribution of pool sizes depends on gene drive carrier frequency, and the sensitivity and specificity of the RDT for each pool size. A maximum likelihood-based approach to designing optimal pooling strategies for given parameter values and subject to given sample size and test kit constraints is described in **Additional file 1**. Here, to evaluate each pooling design, mosquito sampling and pooled testing is simulated *n*_*rep*_ times, and carrier frequency is estimated using a maximum likelihood framework applied to the replicate-level pooled test outcomes. The SD of the estimator, 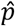, is then derived from the inverse of the observed Fisher information, and pooling designs are ranked in terms of their SD.

We developed an R-Shiny application, “TPP Explorer” (available at https://pverma.shinyapps.io/tpp_explorer), to explore optimal pooling designs to minimize the SD of the estimator, 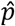, for a given sample size and available number of RDTs. The optimal distribution of pool sizes is highly dependent on gene drive carrier frequency - at high frequencies, larger pools become saturated with positive tests and hence individual tests or smaller pools are more informative; while at low frequencies, larger pools are informative when paired with pools of different sizes. Consider, for instance, a sample size of 100, 20 test kits, a test sensitivity (independent of pool size) of 90% (the ideal value for use case 1), a test specificity (independent of pool size) of 99.95% (the minimally essential value for use case 1), and possible pool sizes of one, two, five and 10. For a carrier frequency of 10%, the optimal pooling distribution is a single test of 10 pooled mosquitoes, 17 tests of five pooled mosquitoes, and two tests of two pooled mosquitoes, which results in an SD of 3.7%. For a carrier frequency of 90%, the optimal pooling distribution is 20 tests of individual mosquitoes, which only makes use of a fifth of the sampled mosquitoes, but results in an SD of 9.7%.

A trade-off exists between the desired precision of the carrier frequency estimate and the number of test kits used; but 20 test kits are generally sufficient to obtain an SD in the vicinity of 10% for the above-listed parameters (a sample size of 100, sensitivity of 90%, and specificity of 99.95%). E.g., for a carrier frequency of 50%, the optimal pooling with 20 test kits results in an SD of 11.6%, which, if desired, can be reduced to 10.0% by increasing the number of test kits to 27.

## 5. Conclusions

Development of an RDT for detecting gene drive mosquitoes will be a critical step toward engaging communities in monitoring activities, enhancing regulatory oversight, and promoting transparency in field trials. Here, we define a TPP for this monitoring tool, establishing minimally essential and ideal criteria to guide test kit development. We consider two use cases: i) detection of a gene drive construct at a location where its presence has not yet been confirmed, and ii) approximate estimation of gene drive carrier frequency, which could be followed up with more extensive laboratory-based assays, when needed. We find that criteria for use case 1 automatically satisfy criteria for use case 2, and hence present these jointly. Field compatibility and ease of use are of primary importance, and the RDT would ideally detect a molecule (DNA, RNA or protein) that would be conserved between the various gene drive implementations under consideration. Test specificity should be very high (at least ≥99.95%), as false positives may create unease, and likely trigger costly follow-up collections and analyses. Test sensitivity should be moderate-to-high (at least ≥60%, and ideally ≥90%). Accurate functioning for pooled testing is ideal, as this would facilitate the use of less test kits in resource-limited settings. An RDT with these characteristics would complement existing molecular surveillance systems, extend monitoring capacity to remote settings, and contribute to building public trust, an essential requirement for the progression of gene drive interventions from laboratory to field. Ultimately, a field-compatible, low-cost RDT would advance safe and ethical implementation of gene drive strategies, and hence contribute to global malaria elimination efforts.

## Availability of data and materials

All codes and source files used to produce the figures in this manuscript are available on GitHub at https://github.com/prateek1verma/tpp-explorer. We have also developed an interactive R-Shiny application called “TPP Explorer,” available at https://pverma.shinyapps.io/tpp_explorer/, which allows users to explore how different test performance characteristics and sampling parameters influence the design and interpretation of diagnostic strategies for gene drive detection. The tool supports flexible investigation of a variety of scenarios, extending beyond the specific examples presented in **Figs. 2 and 3**, including optimal pooling scenarios to minimize SD in the estimator, 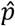, for a given sample size and available number of RDTs. It also allows users to visualize estimated gene drive prevalence across a range of true prevalence values for any pooled test design, with customizable sensitivity and specificity for each pool size.

## Abbreviations

ACT: Artemisinin combination therapy drug
DNA: Deoxyribonucleic acid
ELISA: Enzyme-linked immunosorbent assay
gRNA: Guide ribonucleic acid
IRS: Indoor residual spraying with insecticides
LAMP: Loop-mediated isothermal amplification
LLIN: Long-lasting insecticide-treated net
mRNA: Messenger ribonucleic acid
MRR: Mark-release-recapture
NMCP: National malaria control program
PCR: Polymerase chain reaction
PSC: Pyrethroid spray catch
qPCR: Quantitative polymerase chain reaction
RDT: Rapid diagnostic test for the gene drive construct
RNA: Ribonucleic acid
SOP: Standard operating procedure
SD: Standard deviation
TPP: Target product profile

## Acknowledgements

The authors would like to thank Isabelle Coche, Ana Kormos and Naima Sykes for comments on the manuscript, and Dr. Godrick Oketch for insights to refine the optimal pooling design algorithm.

## Author contributions

PV, SV, CKY, FR, DMM, JK, NW, MS, FT and JMM contributed content and perspectives to the manuscript. SV, CKY and FT drafted the target product profile. PV and JMM performed mathematical modeling. PV, SV, CKY, FR, DMM, JK, NW, MS, FT and JMM reviewed and approved the final manuscript.

## Ethics declarations

### Ethics approval and consent to participate

Not applicable.

### Consent for publication

Not applicable.

### Supplementary information

**Additional file 1**. Optimizing pooled testing for estimation of gene drive carrier frequency.

