## Additional file 1 for "A target product profile for a rapid diagnostic test to monitor mosquito gene drive presence and frequency"

### Additional file 1:
Optimizing pooled testing for estimation of gene drive carrier frequency


Prateek Verma*[1], Sebald Verkuijl*[2], Calvin K. Yee*[2], Filippo Randazzo[3], Damaris Matoke-Muhia[4], Jonathan Kayondo[5], Nikolai Windbichler[2], Michael R. Santos[6], Frédéric Tripet†[7,8], John M. Marshall†[1]

[1]*Divisions of Biostatistics & Epidemiology, School of Public Health, University of California, Berkeley, California, United States of America*
[2]*Department of Life Sciences, Faculty of Natural Sciences, Imperial College London, London, United Kingdom*
[3]*Leverage Science, LLC, Berkeley, California, United States of America*
[4]*Centre for Biotechnology Research and Development, Kenya Medical Research Institute, Nairobi, Kenya*
[5]*Department of Entomology, Uganda Virus Research Institute, Uganda*
[6]*Foundation for the National Institutes of Health, North Bethesda, Maryland, United States of America*
[7]*Swiss Tropical and Public Health Institute, Kreuzstrasse 2, Allschwil, Switzerland*
[8]*University of Basel, Petersplatz 1, Basel, Switzerland*


### 1. Background

During field releases of genetically modified mosquitoes, estimating the prevalence of gene drive carriers will be important to access their spread and long-term stability, particularly as prevalence varies across time and space. Moreover, constraints such as low mosquito abundance, limited trapping capacity, and high diagnostic costs may make it impractical to test each sampled mosquito during their monitoring. Pooled testing through rapid diagnostic kits may offer an alternative to obtain a quick and rough estimate of the gene drive carrier frequency; but introduces greater uncertainty into the estimation process. In this analysis, we aim to identify optimal pooling strategies that could maximize the precision and accuracy of the estimated prevalence while respecting operational constraints on sample size and testing capacity. A central question this analysis seeks to answer is: *Which pooling strategies yield the most reliable estimates of gene drive frequency under practical constraints on sample availability and testing capacity?*


*Equal contribution.

### 2. Modeling framework

We model the sampling process using pooled testing, in which mosquitoes are combined into groups of varying sizes and tested collectively rather than individually. A pooling design is defined by $g$ groups, where each group $i$ is characterized by its pool size $m_i$ (the number of mosquitoes per test) and the number of independent pools $k_i$. Diagnostic performance may vary with pool size, so each group is assigned its own sensitivity $s_i$ and specificity $c_i$. Mosquitoes are assumed to be sampled randomly from the population, and pool outcomes are treated as statistically independent. To assess the performance of a given design, we simulate the entire experiment $n_{\text{rep}}$ times under a fixed true gene drive frequency, generating repeated pooled outcomes for the same design parameters.

#### Pool-level test outcome probability

Let $p$ denote the true gene drive carrier frequency in the population. For a pool of size, $m_i$, the probability that it contains at least one gene drive carrier is:

$$x_{m_i} = 1 - (1 - p)^{m_i}$$

A pool will test positive either because it contains at least one gene drive carrier and the test detects it (true positive), or because of a false positive error. For a diagnostic test with sensitivity, $s_i$, and specificity, $c_i$, of a pool of size, $m_i$, the probability that the test result is positive for the presence of the gene drive carrier is:

$$\theta_i = s_i \cdot x_{m_i} + (1 - c_i) \cdot (1 - x_{m_i})$$

The above expression accounts for both true positives (pools that contain gene drive carriers and are correctly identified) and false positives (pools without gene drive carriers that test positive due to imperfect specificity).

#### Stochastic test outcomes

To evaluate how well each pooling design performs, we simulate the entire experiment $n_{\text{rep}}$ times using the same design parameters and true gene drive frequency. In each replicate, $r$, every group $i$ contributes $k_i$ independent pooled test results.

Assuming statistical independence of pools, the number of positive test results, $y_i^{(r)}$, observed in group $i$ during replicate $r$ follows a binomial distribution with parameters $k_i$ and $\theta_i$, i.e.,

$$y_i^{(r)} \sim \text{Binomial}(k_i, \theta_i).$$

The complete pooling design is defined by vectors of pool sizes, $\vec{m} = (m_1, \ldots, m_g)$, number of pools per group, $\vec{k} = (k_1, \ldots, k_g)$, and pool-level test positive probabilities, $\vec{\theta} = (\theta_1, \ldots, \theta_n)$.

Across all $g$ groups and $n_{\text{rep}}$ replicates, the test outcomes are stored in an $n \times n_{\text{rep}}$ matrix, $\mathbf{Y}$, where each entry $Y_{i,r} = y_i^{(r)}$. This observed outcome matrix, $\mathbf{Y}$, serves as the basis for inference about the underlying gene drive carrier frequency, $p$.

#### Likelihood function

We estimate the gene drive carrier frequency, $p$, using a maximum likelihood framework applied to pooled test outcomes. For the $r$-th replicate, the probability of observing $y_i^{(r)}$ positive pools out of $k_i$ tests in group $i$, given the pool-level probability of testing positive $\theta_i$, is

$$\Pr(y_i^{(r)} \mid p) = \binom{k_i}{y_i^{(r)}} \theta_i^{y_i^{(r)}} (1-\theta_i)^{k_i - y_i^{(r)}}.$$

Assuming independence across replicates and groups, the full likelihood over all $n_{\text{rep}}$ replicates and $g$ groups is

$$L(p) = \prod_{r=1}^{n_{\text{rep}}} \prod_{i=1}^{g} \binom{k_i}{y_i^{(r)}} \theta_i^{y_i^{(r)}} (1-\theta_i)^{k_i - y_i^{(r)}}.$$

Since the binomial coefficients depend only on the observed data, they can be treated as constants with respect to $p$ and dropped from the optimization. Taking logarithms and aggregating outcomes across replicates yields

$$\log L(p) = \sum_{i=1}^{g} \left[ \bar{y}_i \log \theta_i + (K_i - \bar{y}_i) \log(1-\theta_i) \right],$$

where $\bar{y}_i = \sum_{r=1}^{n_{\text{rep}}} y_i^{(r)}$ is the total number of positives observed in group $i$, and $K_i = n_{\text{rep}} \cdot k_i$ is the total number of tests. The maximum likelihood estimate is then obtained as

$$\hat{p} = \underset{p \in (0,1)}{\arg\max} \; \log L(p).$$

#### Boundary cases and implementation details

In practice, direct maximization of the likelihood can be numerically unstable, particularly near the boundaries of the parameter space. To address this, pool-level probabilities $\theta_i$ are clamped away from exact 0 or 1, and optimization is restricted to an interior interval $(10^{-8}, 1-10^{-8})$. The algorithm also incorporates explicit boundary checks: if all pools are negative ($\hat{p} = 0$) or all are positive ($\hat{p} = 1$), these degenerate cases are returned directly. More generally, the log-likelihood is evaluated at both boundaries ($p = 0, 1$) and compared with the interior maximum, with the final estimate chosen as the value of $p$ that achieves the highest likelihood:

$$\hat{p} = \underset{p \in \{0,\, 1,\, p_{\text{int}}\}}{\arg\max} \; \log L(p).$$

These safeguards ensure that the estimator remains computationally stable and statistically well-defined, even in extreme scenarios where observed data provide limited information.

#### Standard deviation from likelihood curvature

Once the MLE $\hat{p}$ is obtained, the standard deviation (SD) is computed from the curvature of the log-likelihood at the maximum:

$$\text{SD}(\hat{p}) = \sqrt{-\, n_{\text{rep}} \,/\, \ell''(\hat{p})},$$

where $\ell''(\hat{p})$ denotes the second derivative of the log-likelihood evaluated at $\hat{p}$. This approach follows standard asymptotic maximum likelihood theory, where the variance of the estimator is approximated by the inverse of the observed Fisher information. In this framework, a sharper curvature corresponds to greater information in the data and hence a more precise estimate of $p$.

#### Optimization criteria and operational constraints

To evaluate the performance of each pooling design, we focused on its ability to provide reliable estimates of the true gene drive frequency, $p$. Practical feasibility was enforced through two constraints: the total number of mosquitoes tested could not exceed $n$, and the number of tests performed was fixed at $K$. Accordingly, only designs satisfying $\sum_{i=1}^{g} m_i k_i \leq n$ and $\sum_{i=1}^{g} k_i = K$ were considered. These conditions reflect operational constraints on mosquito availability and diagnostic capacity in field settings.

For each viable design, we simulated replicate datasets to generate the total number of positive pools observed in each group. From these aggregated outcomes, we estimated the carrier frequency ($\hat{p}$) by maximum likelihood and calculated the standard deviation from the curvature of the log-likelihood at $\hat{p}$, using the observed Fisher information. Designs were then ranked by their SD, with lower values indicating more consistent and precise estimates.

It should be noted that this evaluation captures only sampling variability—how much $\hat{p}$ fluctuates due to random inclusion of mosquitoes in pools. It does not account for uncertainty in the true population value of $p$ itself.

### 3. Simulation strategy

We used simulation to evaluate and compare pooling designs under the operational constraints of mosquito availability and diagnostic capacity. For each candidate design, defined by pool sizes $\vec{m}$ and number of pools $\vec{k}$, we generated $n_{\text{rep}}$ synthetic datasets by simulating binomial outcomes under a fixed true gene drive frequency $p$ and diagnostic parameters $(\vec{s}, \vec{c})$. Within each replicate, the maximum likelihood estimate $\hat{p}$ was obtained, and its standard deviation was derived from the curvature of the log-likelihood function using aggregated outcomes across replicates. Designs were then ranked by their estimated SD, with lower values indicating more precise inference.

The simulation and estimation workflow can be summarized as follows:

**Input:** True frequency $p$; diagnostic parameters $(\vec{s}, \vec{c})$; pool sizes $\vec{m}$; candidate number of pools $\vec{k}$; limits on total mosquitoes $n_{\max}$ and tests $K_{\max}$; number of replicates $n_{\text{rep}}$.
**for** each candidate design $(\vec{m}, \vec{k})$ **do**
  Compute total tests $K = \sum_i k_i$ and mosquitoes used $n = \sum_i m_i k_i$.
  **if** $K = K_{\max}$ and $n \leq n_{\max}$ **then**
    Compute probability that a pool of size $m_i$ contains at least one carrier: $x_{m_i} = 1 - (1-p)^{m_i}$.
    Compute test-level probability of a positive result: $\theta_i = s_i x_{m_i} + (1 - c_i)(1 - x_{m_i})$.
    Simulate pooled outcomes for each group and replicate: $Y_{i,r} \sim \text{Binomial}(k_i, \theta_i)$.
    Aggregate results: $\bar{y}_i = \sum_{r=1}^{n_{\text{rep}}} Y_{i,r}$, with total tests $K_i = n_{\text{rep}} k_i$.
    Evaluate log-likelihood $\log L(p)$ using aggregated outcomes.

Maximize $\log L(p)$ to obtain MLE $\hat{p}$.
Estimate SD of $\hat{p}$ from curvature of $\log L(p)$.
Store design, $\hat{p}$, and SD.
**end if**
**end for**
**Output:** Ranked list of viable designs sorted by ascending SD($\hat{p}$).

#### Prevalence estimator simulations

In real monitoring applications, the true prevalence of the gene drive is not known in advance. While optimal pooling designs can be identified under an assumed true prevalence $p$, their performance may vary substantially across different prevalence values. To assess robustness, and to identify regions where bias or variability increase, we developed an interactive prevalence estimator module within the RShiny TPP Explorer app. This tool visualizes how the estimated carrier frequency $\hat{p}$ behaves across the full range of possible true prevalence $p$, thus complementing the optimal design analysis, performed for a fixed $p$, with a more comprehensive picture of expected performance in practice.

This estimation procedure differs from the likelihood-curvature approach used to rank pooling designs in the following manner: for each candidate $p_{\text{true}}$, replicate pooled datasets are simulated, and the maximum likelihood estimate $\hat{p}$ is recalculated independently for each replicate. The resulting distribution of $\hat{p}$ for all $n_{\text{rep}}$ simulations is then summarized by its mean, a 95% interval obtained through simulation-based resampling, and the scatter of replicate values. In this framework, uncertainty in terms of SD is quantified directly from repeated simulations rather than approximated via the second derivative of the log-likelihood.

This strategy contrasts with the approach used in design optimization, where thousands of candidate configurations must be compared efficiently. In that setting, the SD of $\hat{p}$ is estimated from the curvature of the likelihood surface to reduce computational cost. By comparison, the prevalence estimator focuses on a single user-specified design and can therefore employ simulation-based resampling to provide a more detailed and transparent characterization of estimator bias and variability across a wide range of true prevalence values. An interactive implementation of these simulations (TPP Explorer) is available at https://pverma.shinyapps.io/tpp_explorer/.
